# A chromosome-level genome assembly of a model conifer plant, the Japanese cedar, *Cryptomeria japonica* D. Don

**DOI:** 10.1101/2023.02.24.529822

**Authors:** Takeshi Fujino, Yamaguchi Katsushi, Toshiyuki T Yokoyama, Toshiya Hamanaka, Yoritaka Harazono, Hiroaki Kamada, Wataru Kobayashi, Tokuko Ujino-Ihara, Kentaro Uchiyama, Asako Matsumoto, Ayako Izuno, Yoshihiko Tsumura, Atsushi Toyoda, Shuji Shigenobu, Yoshinari Moriguchi, Saneyoshi Ueno, Masahiro Kasahara

## Abstract

Japanese cedar (*Cryptomeria japonica* D. Don) is the most important Japanese forest tree, occupying about 44% of artificial forests in Japan, and planted in East Asia, Azores Archipelago, and some islands in the Indian Ocean. Although the huge genome of the species (ca. 11 Gb) with abundant repeat elements might have been an obstacle for genetic analysis, the species is easily propagated by cutting, flowered by plant hormones like gibberellic acid, transformed by agrobacterium, and edited by CRISPR/Cas9. These characteristics of *C. japonica* are preferable to make the species a model conifer for which reference genome sequences are necessary. In this study, we report the first chromosome-level assembly for *C. japonica* (2n = 22) using a third generation selfed progeny with an estimated homozygosity of 0.96. Young leaf tissue was used to extract high-molecular-weight DNA (>50 kb) for HiFi PacBio long read sequencing and to construct Hi-C/Omni-C library for Illumina short read sequencing. Using the 29× and 26× genome coverage of HiFi and Illumina reads, respectively, de novo assembly resulted in 2,650 contigs (9.1 Gb in total) with N50 contig size of 12.0 Mb. The Hi-C analysis mapped 97% of the nucleotides on the 11 chromosomes. The assembly was verified by comparing with a consensus linkage map of 7,785 markers. The BUSCO analysis confirmed ~91% of conserved genes. Annotations of genes, repeat elements and synteny with other Cupressaceae and Pinaceae species were performed, providing fundamental resources for genomic research of conifers.

## Introduction

Conifers are known to possess large genomes (6.5 to 37 Gbp) and an abundance of repetitive sequences in comparison to the genomes of animals such as humans [1–6]. As a result, the cost of determining the genome sequence of a conifer used to be quite high. In addition, creating genetic maps for these species takes a long time due to the long generation cycle of conifer, leading to a limited application of genetic methods (e.g., genetic markers, transgenesis, gene editing) in conifers; efforts at breeding via molecular biological techniques based on both genetic maps and genome sequences are limited only to a few species. Therefore, there is a pressing need for constructing a chromosome-level genome assembly of a conifer species for efficient molecular breeding, although assembling a conifer genome with abundant repeat sequences was challenging. Such a genome assembly would facilitate breeding elite trees with a variety of favorable quantitative and qualitative traits, including rapid growth rate, resistance to pathogens, and desired timber quality [7,8].

With the advent of accurate long read sequencing methods such as PacBio HiFi reads or Oxford Nanopore kit v14 has significantly decreased the cost of determining large genomes [4,9,10], leading us to believe that now is an opportune time to establish a high-accuracy, high-contiguity model conifer genome at the chromosome scale with comprehensive gene annotation. The chromosome-level assembly of a conifer genome and its associated high-density genetic map and gene annotation would facilitate the rapid identification of the relationship between various traits and genes (or mutations) on a molecular basis in a systematic manner.

Japanese cedar or sugi (*Cryptomeria japonica* D. Don), which has a large genome size (about 11 Gb estimated from DNA content (22.09pg/2C; [11]), is an allogamous and monoecious conifer, naturally distributed from Aomori Prefecture to Yakushima Island in Japan [12]. *C. japonica* is a forestry species that covers the largest area of any planted trees in Japan, making it the most important species in Japan’s tree industry. In particular, it was extensively planted for timber production in the 1960s to support the country’s economic growth. Currently, the species occupies nearly 4.5 million hectares, accounting for 44% of all artificial forests in Japan [13]. The proliferation of artificial forests of *C. japonica* has led to a widespread prevalence of pollinosis in Japan, with a reported prevalence of 38.8% and corresponding major social issues [14]. There is therefore a pressing need to reduce the amount of *C. japonica* pollen dispersed by renewing them with male-sterile trees of *C. japonica*. Indeed, sequence-based markers for male sterility were established in this line of research [15]. These sequence-based markers allow for the easier identification of wild trees (for breeding) with a male-sterility allele; the estimated frequency of trees with male-sterile alleles in breeding materials is approximately 1% [16,17]. Had there been a chromosome-level assembly, developing genetic markers for marker-assisted selection would have been significantly facilitated. Given the industrial and social needs for the genome of *C. japonica*, we initiated PacBio HiFi sequencing of the genome of *C. japonica* with the goal of establishing a chromosome-level assembly covering over 95% of the nucleotides on the eleven chromosomes of the *C. japonica*.

Here we present a highly contiguous, chromosome-level assembly of *C. japonica* using a combination of HiFi sequencing, Hi-C scaffolding and a high-density genetic map. Our chromosome-level assembly of *C. japonica* genome is the first genome sequence in a conifer species with high sequence contiguity (N50 contig >12 Mb), with the majority of nucleotides mapped to the chromosomes (>97% assigned to the 11 chromosomes (2n=22)), and with validation through high-density genetic map (7,781 sequence-based markers; Supplementary Table 6). Genes are annotated using existing and newly sequenced RNA-seq data, ESTs in public databases, Iso-seq previously reported in literature, and homology to proteins in other species, resulting in an unprecedented high completeness score (91.4%) of BUSCO [18]. This newly sequenced genome will serve as a reliable foundation for breeding and studying other conifer species as well; for instance, it is valuable for identifying loci associated with tolerance to pathogens, faster growth, adaptation to various environments, and the evolution of conifer genomes and retrotransposons.

## Results

### Breeding Homozygous Trees

Assembling highly heterozygous genomes had long been known to be challenging [3,19–21], especially when we initiated our sequencing project of *C. Japonica* prior to the advent of PacBio HiFi sequencing. To reduce the heterozygous regions of the genome as much as possible, we attempted to use inbred *C. japonica* trees. However, we were unable to obtain a fully homozygous genome like inbred lines seen for rice or maize. As the self-fertilization of more than four times was not accomplished, we decided to use ‘Kunisaki-3 S3-3’, a third-generation inbred tree with the lowest heterozygosity among the selfed progenies, for genome sequencing. The heterozygosity rate of the selected tree was confirmed to be approximately 0.96 through a SNPType assay (Fluidigm) with 158 SNP markers (Supplementary Table 1), consistent with the results of GenomeScope analysis, which will be presented later.

### Genome Sequencing

#### Genome DNA preparation from Young Leaves

Young leaves in April and May were sampled from ‘Kunisaki-3 S3-3’. To prepare high molecular weight (HMW) DNA from *C. japonica*, the sample ground into powder in liquid nitrogen was mixed with 50 ml of a pre-wash buffer [50 mM Tris-HCl (pH 8.0), 10 mM EDTA, 0.2% 2-ME, 0.3% TritonX] and incubated at room temperature for 1 hour with gentle agitation. This pre-wash step serves to reduce the viscosity of the extracted DNA solution. Following centrifugation at 9,000 × *g* for 20 minutes at 4°C, the pellet sample was snap-frozen. Fifty ml of frozen powdered QIAGEN G2 buffer (Qiagen, Netherlands, Cat #1014636), which was produced by spraying the buffer into liquid nitrogen in a glass beaker, was added to the sample and mixed quickly. Upon allowing the mixture to thaw in a tube, 30 ul of RNaseA (QIAGEN, Cat #19101) and 2.5 ml of Proteinase K (QIAGEN, Cat #1019499) were added, and the sample was incubated at room temperature for 2 hours without agitation. The sample was centrifuged at 9,000 × *g* for 20 minutes at 4°C, and the supernatant was subjected to DNA extraction using two QIAGEN Genomic-tip 100/G columns (Cat # 10243). The genomic DNA was eluted with 5 ml of Buffer QF, precipitated with isopropyl alcohol, and dissolved with 600 ul of TE at 4°C for a week, which yielded 247 μg of DNA (412 ng/μl). We repeated this procedure to obtain another batch of HMW genome DNA from the young leaves of the same tree, yielding 356 μg of DNA (594 ng/μl). The size distribution of the HMW DNA was evaluated by pulsed-field gel electrophoresis (PFGE). Briefly, 20 ng of HMW DNA was run on a 1% agarose gel (Seakem Gold Agarose, Lonza, Rockland, ME, USA, Cat #50150) in 0.5x TBE with the BioRad CHEF Mapper system (BioRad, Hercules, CA, USA, Cat #M1703650) for 15 hours, and the gel was stained with SYBR Gold dye (Thermo Fisher Scientific, Waltham, MA, USA, Cat #11494). Lambda ladder, *Saccharomyces cerevisiae* genome, and 5 kbp ladder (BioRad,Cat #170–3624) were used as standards. These results demonstrated that the HMW DNA had an approximate mean size of 50 kb. DNA concentration was measured using a Qubit fluorometer (Thermo Fisher Scientific, Cat #Q32866) and the Qubit™ dsDNA BR Assay Kit (Thermo Fisher Scientific, Cat #Q32850). The quality of the extracted DNA was evaluated by analyzing the absorbance spectra with Nanodrop ND-2000C (Thermo Fisher Scientific).

#### NGS Library preparation and sequencing

##### PCR-free Illumina Read Sequencing

For PCR-free Illumina library preparation, we used 6 ug of the genomic DNA (gDNA) extracted using the QIAGEN Genomic-tip 100/G columns following the method described in the previous section. The gDNA was fragmented into 200-800 bp fragments with a peak at 550 bp using the Covaris Focused-ultrasonicator S2 system with the following settings: intensity=4; duty cycle=10%; cycles per burst=200; time=55 sec; temperature=7°C. Subsequently, ~400-bp fragments were size-selected by using BluePippin and 2% EF gel in the tight mode setting, followed by concentration with AMpure XP (1.8x vol.), yielding 226 ug of DNA with a sharp peak at 410 bp as evaluated with Agilent 2100 Bioanalyzer. A library for whole genome sequencing was prepared using the TruSeq DNA PCR-Free Library Prep Kit (Illumina) according to the manufacturer’s instructions. The concentration of the library was quantified by KAPA Library Quantification Kit and Applied Biosystems 7500 Real-Time PCR Systems (Applied Biosystems). The Illumina library was sequenced using the HiSeq X platform (Illumina) at Macrogen Japan (Tokyo, Japan) with the 2×151 bp paired-end sequencing protocol. The total number of the raw Illumina paired-end reads was 734439281.

##### HiFi Long Read Sequencing

HMW genome DNA obtained through the procedures described above was shared to an average target size of 20-30 kbp via centrifugation in a Covaris g-Tube (cat# 520079) at 2800 - 4800 rpm for 2 min using a microcentrifuge (himac CG15RX). SMRTbell libraries for sequencing on the PacBio Sequel platform were constructed according to the manufacturer’s recommended protocol (Preparing HiFi SMRTbell Libraries using SMRTbell Express Template Prep Kit2.0 PN101-853-100 Version 03 (January 2020) or Version 04 (April 2021)) with minor modifications. We altered the condition of genome shearing with the g-Tube and the size selection of the library with the BluePippin system (Sage Science,. MA,USA.) with the aim of constructing libraries with longer inserts. Specifically, we constructed seven SMRTbell libraries with various settings of centrifugation speed, including 4800 rpm, 3800 rpm, 4300 rpm, 4800 rpm, 3300 rpm, 2800 rpm, and 2800 rpm, yielding libraries with average sizes of 20 kb, 20 kb, 18 kb, 16 kb, 22 kb, 25 kb, and 25 kb, respectively. These insert size distributions were determined by PFGE with the BioRad CHEF Mapper system as previously performed for HMW genome evaluation (see above). These SMRTbell libraries were sequenced in 1 SMRT cell (movie time of 20 hours) on a Sequel instrument at the NIBB Functional Genomics Facility and 10 cells (movie time of 30 hours) on a Sequel II instrument at the National Institute of Genetics, ROIS, Japan (Supplementary Table 3).

##### Hi-C Sequencing

The Omni-C library was prepared using the Dovetail Omni-C Kit (Dovetail cat# 21005) according to the manufacturer’s protocol (Omni-C Protocol Non-mammal v1.0 for Plants) with modifications. Due to our inability to obtain chromatin through the standard in-place fixation procedure from leaves, we employed a third-party protocol [High Molecular Weight DNA Extraction from Recalcitrant Plant Species for Third Generation Sequencing by Workman; dx.doi.org/10.17504/protocols.io.4vbgw2n], to isolate nuclei from leaf samples, after which we proceeded to nuclease digestion. The ends of chromatins were polished and ligated to a biotinylated bridge adapter prior to proximity ligation of adapter-containing ends. Following proximity ligation, crosslinks were reversed, and DNA was purified from proteins and size-selected. Purified DNA was subjected to end-repair, ligated to an adapter, and treated with USER Enzyme. Four sequencing libraries were generated using Illumina-compatible adapters, and biotin-containing fragments were isolated using streptavidin beads before PCR enrichment of the library. These libraries were sequenced on two lanes of the Illumina HiSeq X platform, yielding 470 million 2 × 151 bp read pairs.

### Transcriptome Sequencing

In order to perform gene annotation for *C. japonica*, we carried out RNA-Seq for several tissues: root, corn, female flower bud, and seeds in corn (Supplementary Table 2). Each tissue sample (50 to 100 mg) was disrupted under liquid nitrogen, and the tissue powder was emulsified by 600 μL of CTAB extraction buffer (Supplementary Table 2) and incubated for 10 min. at 65 °C. After centrifugation for 5 min. at 15,000 rpm at 4 °C, the supernatant (400 μL) was mixed with 200 μL of Lysis Buffer of the Maxwell® RSC Plant RNA Kit (Promega). RNA was then extracted by the Maxwell® instrument (Promega) according to the manufacturer’s instructions. The extracted RNA was sent to a sequencing company (Macrogen) to construct sequencing libraries and sequenced on NOVASeq6000 (Illumina) in 2 × 101 bp paired-end form. Publicized RNA-Seq data (Supplementary Table 2) [22,23] were also used in the current study.

### Genome Assembly

#### Assembling PacBio HiFi reads

The collective statistics of the obtained PacBio HiFi reads are displayed in Table 1. Through the use of FastQC [24], we discovered that the first 12+ bases occasionally contained adapter sequences that were not filtered by the official PacBio pipeline (Supplementary Figure 1). After cutting the first 20 base pairs, the base frequency bias due to the adapters disappeared (Supplementary Figure 2). To see how the remaining adapters affect the assembly statistics, we generated three sets of the HiFi reads by trimming the first 0, 20, 30 base pairs using fastp ver 0.20.0 [25]. Then we assembled the three sets of the PacBio HiFi reads using hifiasm 0.15.5-r350 [26] and Flye 2.8.3 [27]. However, the Flye assembly jobs did not complete due to out of memory (> 300 GB), and we estimated that even with infinite memory the job would take more than 62 days, which is the maximum computation time for our shared computing environments. Hifiasm completed in two days, so we used the result of the Hifiasm (Supplementary Table 2). We selected the assembly with the trim size = 20 bp because it has the longest N50 contig size and the other metrics are largely similar to the other assemblies (Supplementary Table 4). The final contigs had a N50 contig size of 12 Mb, which is significantly larger than those for previously published conifer genomes such as [4]. The total contig size is 9.07 Gbp, which is close to the estimated genome size. The statistics for the final contigs are presented in Table 2 below.

**Table 1:**
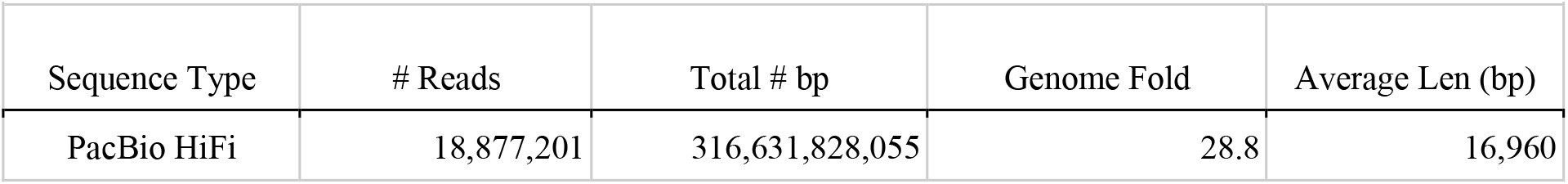
Sequencing Statistics.

| **Table 1: Sequencing Statistics.** |  |  |  |  |
| --- | --- | --- | --- | --- |
| Sequence Type | # Reads | Total # bp | Genome Fold | Average Len (bp) |
| PacBio HiFi | 18,877,201 | 316,631,828,055 | 28.8 | 16,960 |

**Table 2:** Assembly Statistics.

| **Table 2: Assembly Statistics.** |  |  |  |  |  |
| --- | --- | --- | --- | --- | --- |
| Assembler | N50 Contig Size (bp) | L50 | Total Count | Total Length (bp) | Max Contig Size (bp) |
| HiFiasm | 12,036,660 | 217 | 2,651 | 9,068,249,119 | 88,939,144 |

#### Hi-C Scaffolding

We utilized Juicer v1.6 [28] and BWA v0.7.17-r1188 [29,30] to align Hi-C reads to the contigs, subsequently utilizing the 3D-DNA pipeline v180922 [28] to correct misjoins, order, and orient the contigs into scaffolds. We subsequently employed Juicebox Assembly Tools v2.13.07 for manual review of the scaffolds, making a few necessary corrections for obvious misassemblies. The 3D-DNA pipeline was subsequently utilized to rescaffold the final chromosome-length scaffolds, with all tools being executed using default parameters. The largest 11 scaffolds, which correspond to the 11 chromosomes of *C. japonica*, contained 97.6% of the nucleotides (Figure 1; Table 3; Supplementary Figure 3).

**Figure 1:**
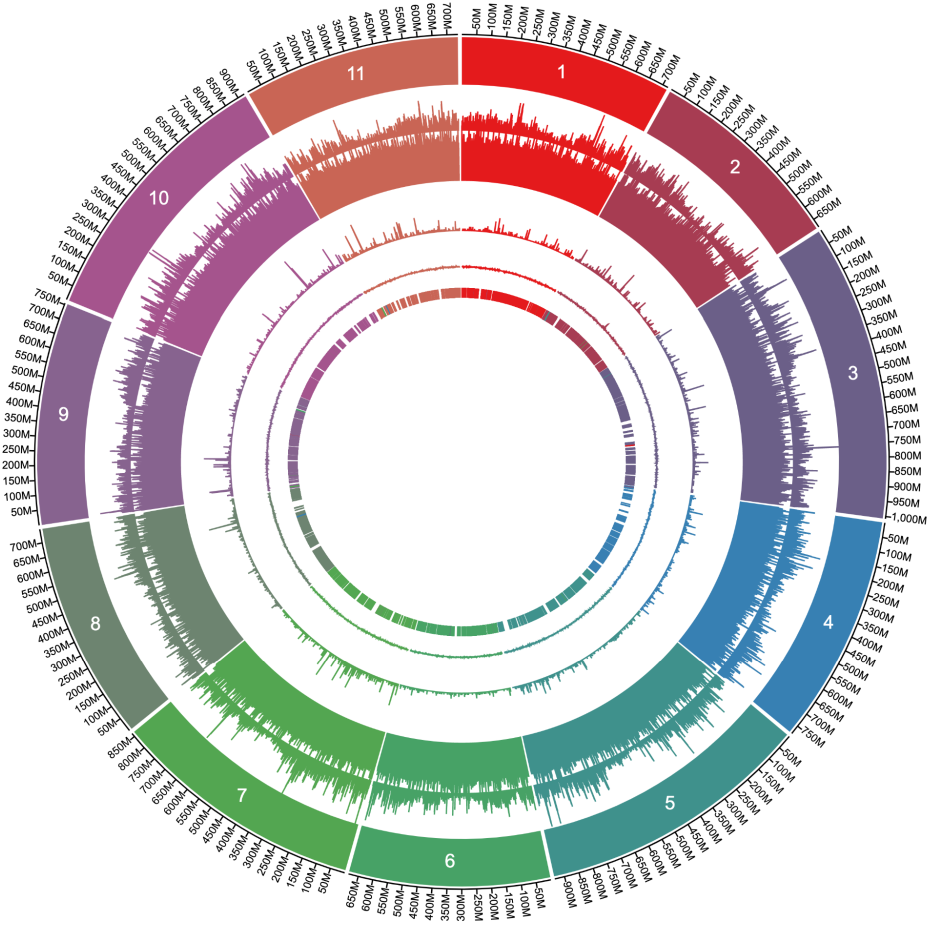
Overview of the 11 chromosomes of the *C. japonica* genome. Rings represent (from outside) chromosomes, gene density, repeat sequence %, N% (gaps between contigs), GC%, and genetic markers (described later). The genome is highly repetitive across the genome.

**Table 3:** Assembly Statistics after Hi-C Scaffolding.

| Table 3: Assembly Statistics after Hi-C Scaffolding. |  |  |  |  |
| --- | --- | --- | --- | --- |
| Top 11 Scaffolds' shows the percentage of the nucleotides on the 11 chromosomes of the genome. |  |  |  |  |
| N50 Scaffold Size (bp) | # Scaffolds | Total Length (bp) | Max Scaffold (bp) | Top 11 Scaffolds (%) |
| 754,841,075 | 2,699 | 9,046,663,964 | 1,007,622,168 | 97.6 |

#### Post-processing of the Hi-C scaffolds

Utilizing NCBI-BLAST [31] with the parameters specified in VecScreen [32], we screened for contaminants such as PacBio adapters within the scaffolds and subsequently masked them as ‘N’s. Any scaffold whose entire sequence was masked was subsequently removed.

To identify scaffolds corresponding to the chloroplast, we employed minimap2 [33] with default parameters to align an existing reference chloroplast sequence (acc: AP009377) against the scaffolds. Subsequent manual review removed scaffolds that largely aligned with the entire reference sequence. A single representative chloroplast sequence was then selected from the genome, reverse complemented and rotated to better align with the existing reference sequence, the annotation of which was transferred to the new sequence using GeSeq [34]. While we attempted to locate mitochondrial sequences as well, we were unable to do so reliably due to the nature of complex mitochondrial genome structure of conifers [35] and thus, potential mitochondrial sequences remain as ordinary scaffolds. The statistics of the final scaffolds are presented below.

### Genome Assembly Validation by Genetic Map

Linkage maps were constructed from four existing mapping families (F1O7, S1-2, S5HK7, and S8HK5) [36] using the LPmerge program [37] (Supplementary Table 6). The sequence-based markers in the merged genetic map are aligned to the scaffolds using minimap2 [33] with the options -c -k 9. The majority of the markers were consistent with the scaffolds (Supplementary Figure 4). As the genetic map was developed independently of the genome assembly, the high level of consistency between the Hi-C scaffolds and the genetic map attests to the high accuracy of the Hi-C scaffolds in terms of minimizing global misjoins. Markers that mapped multiple times may not be consistent and minor inconsistencies resulting from ambiguities during the merging of the four independent genetic maps were also noted (Supplementary Figure 4). The mapped markers are well scattered so any single locus trait would be easily mapped (Figure 1).

It is worth noting that there are long regions where recombination is suppressed in the middle of the chromosomes. We presume that these regions correspond to centromeres. The gene density decreases towards these putative centromeres (Figure 1).

### Identifying Repeat Elements

We ran RepeatModeler 2.0.3 [38] to identify repeat elements in the contigs, yielding a repeat library (Table 4) that was subsequently input to RepeatMasker 4.1.2-p1 [2] for identifying repeat elements in the genome. We used the most sensitive preset, -s option. We used Cross_match [39] for the search engine of RepeatMasker because it is the most sensitive engine. As is common in other conifer species, the *C. japonica* genome is abundant in repeat elements, with 83.6% of the genome identified as such (Table 5). These repeat elements are evenly scattered across the genome, showing no observable bias on the chromosome location (Figure 1). The majority of ‘Unknown’ and ‘LTR/Unknown’ (included in ‘Other LTR’ in Table 5) presumably consists of fragmented copies of *Gypsy* or Copia superfamilies not precisely identified by RepeatModeler2.

**Table 4:** Statistics of the Output of RepeatModeler2.

| # sequences | Total Length | Min. Length | Avg. Length | Max. Length | N50 |
| --- | --- | --- | --- | --- | --- |
| 4,089 | 10,847,836 | 34 | 2,653 | 39,182 | 5,201 |

**Table 5:**
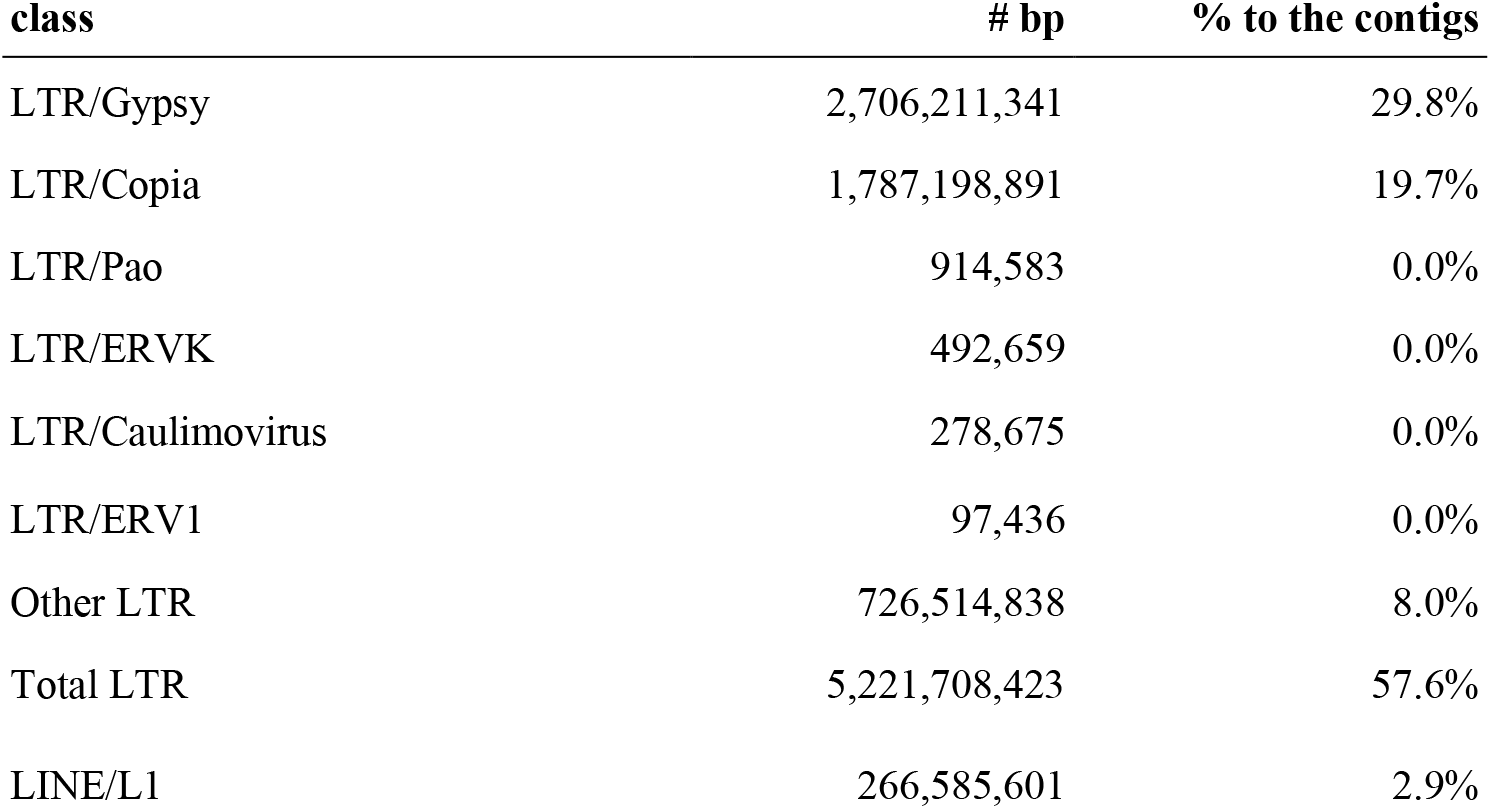
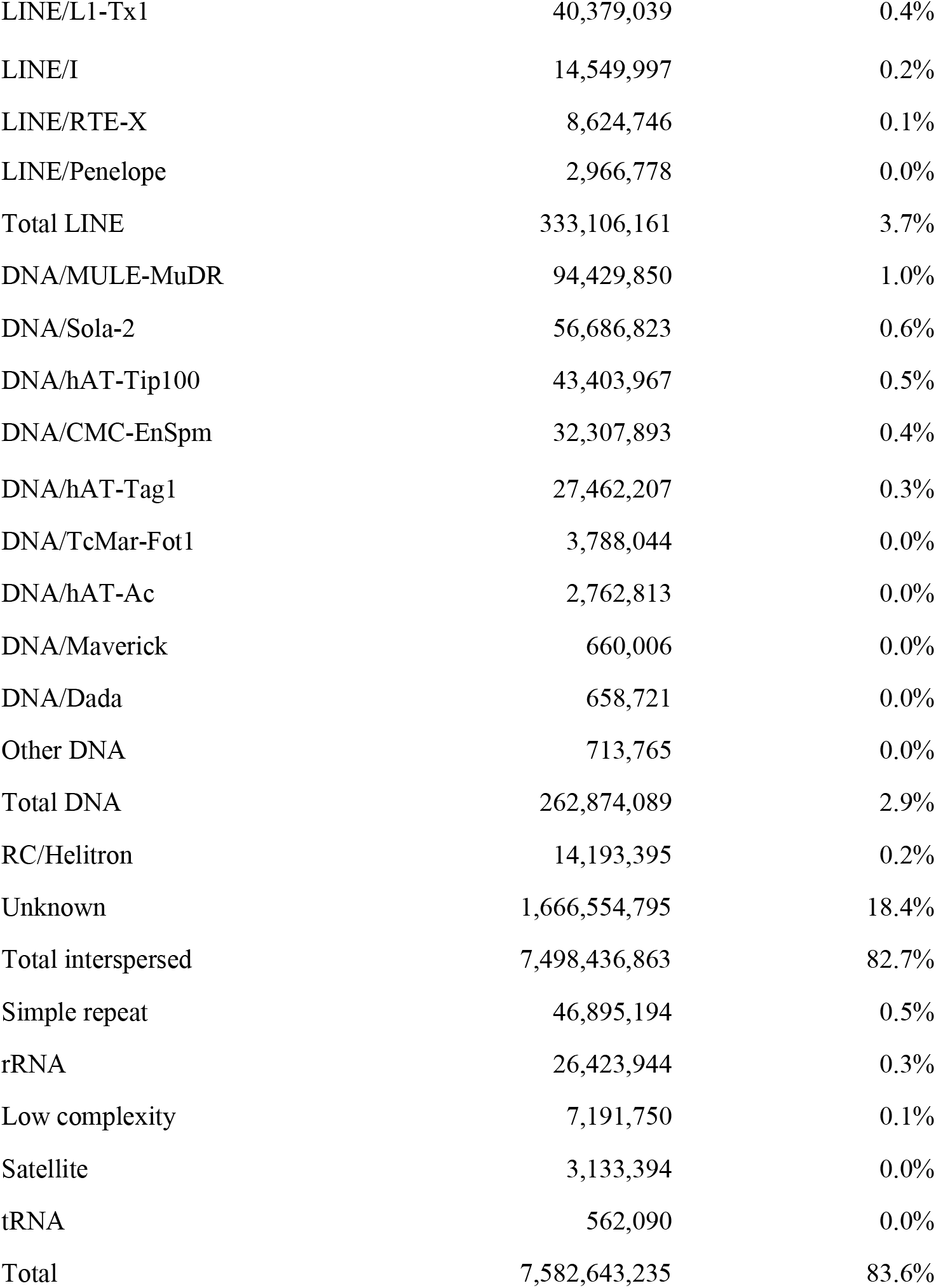
Breakdown of Repeat Elements. Unknown and LTR/Unknown are presumably fragmented LTR/Gypsy and LTR/Copia which RepeatModeler failed to identify the element category.

### Gene and Functional Annotation

We used BRAKER 2.1.6 [40] to predict two sets of genes, one based on RNA-seq data and the other based on homology with protein databases. For the former prediction set, the RNA-seq alignments were used for training the gene model, while the homology-based prediction was conducted using the proteins at the Viridiplantae level of OrthoDB v10 [41–43]. We assembled the RNA-seq data *de novo* using Trinity 2.13.2 [44]. In addition, we performed reference-based transcriptome assembly using StringTie 2.1.7 [45] with the RNA-seq data, ESTs in GenBank/EMBL/DDBJ [22] and Iso-seq reads previously reported in [23]. Subsequently, the *de novo* transcriptome assembly, the ESTs, the Iso-seq reads and the gene structures produced by StringTie were input into the alignment assembly pipeline of PASA 2.5.2 [46,47] to generate an integrated gene structure on the contigs. Finally, EVidenceModeler 1.1.1 [47] was employed to integrate all the gene sets from the BRAKER pipeline and the PASA alignment assembly pipeline. EVidenceModeler output a total of 152,527 genes, which we refer to as *a permissive gene set*, as it contains many false positive genes.

As the permissive gene set contains many false positives, we created another gene set, referred to as *the standard gene set*, for general analysis and databases. To do so, we first removed genes from the permissive gene set that overlap 95% of their length with repeat elements, as these genes are typically proteins almost entirely within transposable elements. We then eliminated genes that lack both at least 5% overlapping length with the PASA transcripts and homology with the proteins in at least one of SwissProt [48], NCBI RefSeq [49], and Viridiplantae of OrthoDB v10 [42] (defined by any statistically significant hits using the default parameter of DIAMOND [50]) are eliminated. In other words, we eliminated predicted genes that lack both expression evidence and homology to other species. In all, the remaining gene set, which we call the standard gene set, contained 55,246 genes. We confirmed that throughout these filtering the BUSCO scores do not change significantly (data not shown). Supplementary Table 7 shows for each gene if it is in the standard gene set, if it has an overlap with the PASA transcripts (RNA-seq evidence), if it has homology to proteins in other species (homology evidence), if it is almost entirely contained in the repeat elements, and if it contains a part of transposable elements (identified by TEsorter ver 1.3 [51] with an options, -db rexdb-plant -dp2 and with Viridiplantae v3.0 of REXdb).

EnTAP 0.10.8 [52] was used for functional annotation with default parameters and following databases: UniProtKB release 2022_05 [48] and NCBI RefSeq plant proteins release 215 [49]. The descriptions of the top protein hit were transferred to the gene descriptions when EnTAP found it informative. When the hit was not informative, we transferred the EggNOG [53,54] description when available.

### Evaluation of Gene Annotation

The gene prediction and annotation for the genome was first done on the contigs and then were lifted over to the chromosome-level assembly. We here evaluated the quality and completeness of our gene annotation using BUSCO v5.3.0 [18,55,56]. As expected from the high contiguity and high accuracy of the HiFi assembly, the BUSCO score for the standard gene set showed very high (> 90%) completeness for the conserved gene set (embryophyta_odb10; see Table 6). Approximately 5% of the duplicated genes (Complete duplicated) may account for the heterozygosity between the two haplotypes (4%) that remained even through three rounds of selfing (see the *Breeding Homozygous Trees* section).

**Table 6:**
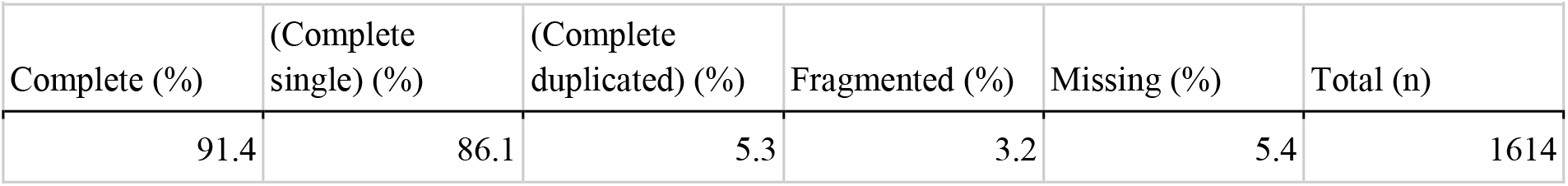
BUSCO Assessment of the Assembly.

| **Table 6: BUSCO Assessment of the Assembly** |  |  |  |  |  |
| --- | --- | --- | --- | --- | --- |
| Complete (%) | (Complete single) (%) | (Complete duplicated) (%) | Fragmented (%) | Missing (%) | Total (n) |
| 91.4 | 86.1 | 5.3 | 3.2 | 5.4 | 1614 |

To our knowledge, the complete gene rate (91.4%) of the *C. japonica* genome is the highest among conifer species reported to have their genomes sequenced, demonstrating the potential of *C. japonica* as a model conifer species.

### Evolution of Repeat Elements

Assembling genomes with abundant repeat elements used to yield contigs with less contiguity (i.e., orders of magnitude smaller N50 contig size) when PacBio HiFi reads were not used for genome assembly. Such genome assemblies tended to selectively miss repeat elements, likely having led to inaccurate analysis of repeat elements. The N50 contig size of our *C. japonica* genome assembly is over 10 Mb; the assembly should contain more repeat elements then ever. This gives us an opportunity to analyze more accurately how repeat elements evolved.

With this in mind, we analyzed the ages of repeat elements in the *C. japonica* genome. We largely followed Ma et al [57]. Briefly, (1) we identified the locations of LTR elements by running EDTA [58], (2) aligned the two LTR copies for each identified LTR retrotransposon; the two copies have exactly the same sequence when it was inserted, and therefore the discrepancies between the two LTR copies can serve as a molecular clock, (3) using the mutation rate of 0.59×10^−9^ /site/year previously reported for *C. japonica* [59], the number of mutations between the two LTR copies is converted to the age of the inserted LTR, and (4) we created the histogram of the age distribution of the LTR retrotransposons for each of *Gypsy* and *Copia* superfamilies (Figure 2). To our knowledge, this is the first description of the age distribution of LTR retrotransposons for conifer genomes with N50 contig size of over a megabase.

**Figure 2:**
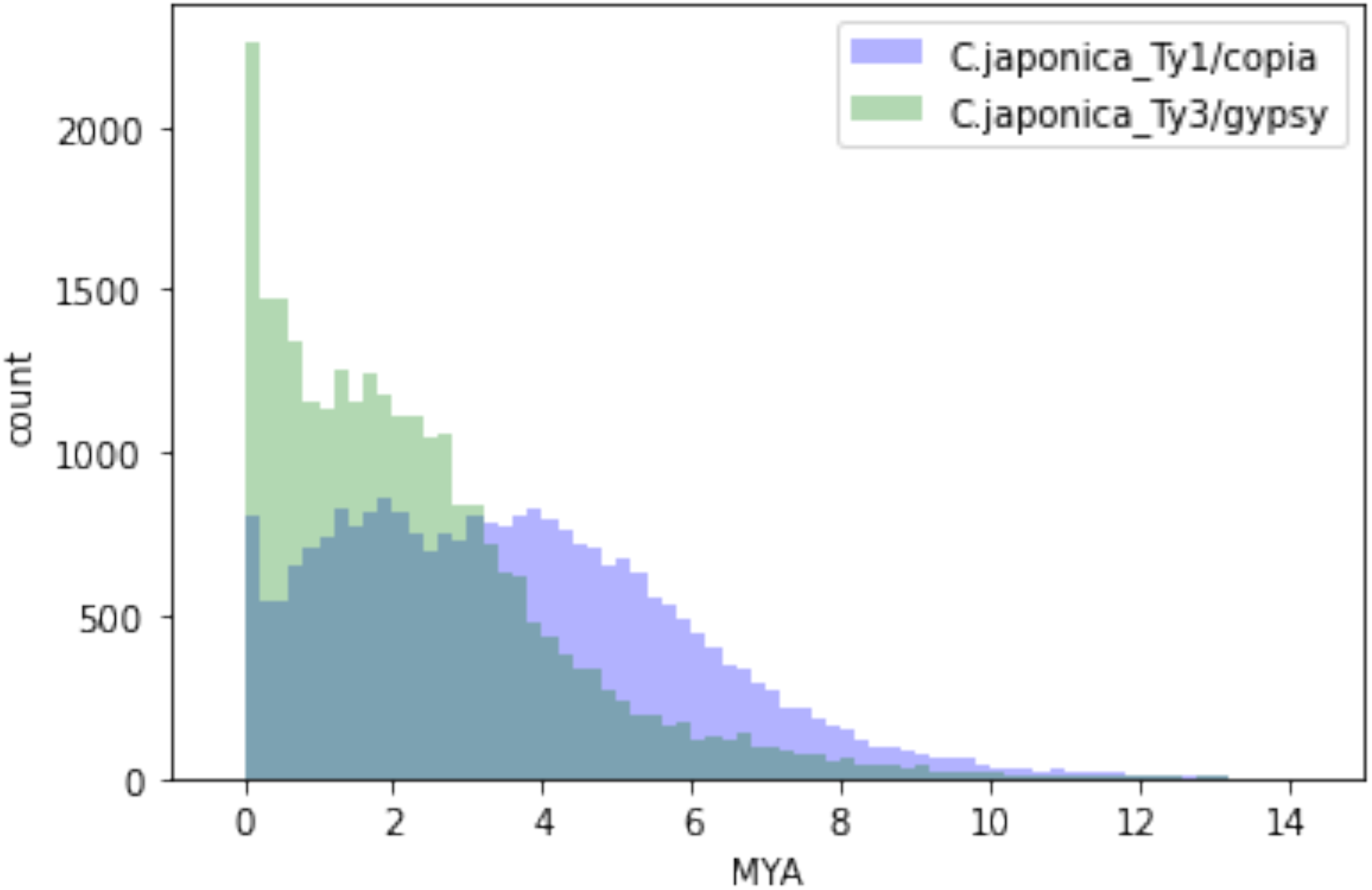
Age distribution of Copia and Gypsy in the *C. japonica* genome. The x-axis is million years old and the y-axis is the number of their occurrences in the genome.

### Genome Browser at ForestGEN

The genome sequence of *C. japonica* can be explored from ForestGEN (FORest EST and GENome database), where JBrowse [60] displays the genomic sequences and gene annotations (https://forestgen.ffpri.go.jp/en/index.html). ForestGEN also has database functions which enable users to query the functional annotation of the predicted genes.

## Discussions

We assembled the *C. japonica* genome sequence with a very high degree of continuity (N50 contig: 12.0 Mb) and were able to map the majority (> 97%) of the nucleotide sequences to all of the 11 chromosomes using the Hi-C scaffolding and the genetic map. We were also able to validate the correctness of the chromosome-level assembly by comparing it with the high-density genetic mapping. This allowed us to establish a genetic information foundation that can be used to develop selecting markers for *C. japonica* breeding in a shorter period of time using a systematic method. In addition, we annotated about 50,000 genes in the *C. japonica* genome sequence and achieved the highest BUSCO score (91.4% complete) among conifers with chromosome-level assembly, to our knowledge. These genome resources thereby reinforce the position of *C. japonica* as a model conifer tree.

*C. japonica* can be used as a model conifer for several reasons. The species is easily propagated by cutting [[61,62], flowered by plant hormones like gibberellic acid [63], transformed by agrobacterium [64], and can be edited by CRISPR/Cas9 [65]. This study aims at providing the high-quality, chromosome-level genome sequence of the *C. japonica*.

One of such breeding efforts is the development of male-steriile trees of *C. japonica* for ameliorating the prevalence of pollinosis in Japan. Since the discovery of male-sterile tree in 1992 [66], breeding of male-sterile trees was initiated including screening of new male-sterile trees in the field, because such variety is expected to reduce airborne pollen dramatically. Four Mendelian inherited male sterility loci (*MS1–MS4*) are known [36], but their exact sequences were not known and the candidate genomic regions for them were too huge to sequence by using tiling clones such as Bacterial Artificial Chromosomes due to the huge genome size.

Sequencing the whole genome of *C. japonica* and narrowing the candidate genomic regions of each of the *MS1–MS4* loci using genetic markers was considered to be the fastest way to obtain the sequence-based markers of *MS1* to *MS4*. Indeed, our chromosome-level assembly of the *C. japonica* genome, which was enabled by the power of long read DNA sequencers and the Hi-C scaffolding, contributed to successfully identify the candidate genes of *MS1* and *MS4* [15,67], having demonstrated the usefulness of our chromosome-level genome sequences.

## Data Availability

The raw PCR-free Illumina reads were deposited in the Sequence Read Archive database of the DNA Data Bank of Japan (DDBJ) under accession number DRX412221, the Omni-C Illumina reads under DRX412222, the PacBio HiFi reads under DRX409629, the assembly and gene annotation under BSEH01000001-BSEH01002698. The assembly, gene annotation, genetic map, and BLAST search are available in the ForestGEN database (https://forestgen.ffpri.go.jp/en/index.html).

## Abbreviations

BAC: Bacterial Artificial Chromosomes
DDBJ: DNA Data Bank of Japan
FFPRI: Forestry and Forest Products Research Institute
HMW: High Molecular Weight
PCR: Polymerase Chain Reaction
ROIS: Research Organization of Information and Systems

## Acknowledgements

This work has been supported in part by FFPRI Grant (#201406, #201421, #201906), JSPS KAKENHI Grant Number JP16H06279 (PAGS), and JP20H03239, Bio-oriented Technology Research Advancement Institution (BRAIN) Grant (JPJ007097; Project ID 28013B), and NIBB Collaborative Research Program (15-829, 16-403, 17-405, 18-408, 19-420, 20-428, 21-302, and 22NIBB402). The super-computing resource was provided in part by Human Genome Center (the University of Tokyo), in part by NIG supercomputer at ROIS National Institute of Genetics, in part by the SuperComputer System, Institute for Chemical Research, Kyoto University, in part by the Data Integration and Analysis Facility, National Institute for Basic Biology, and in part by the supercomputer of AFFRIT, MAFF, Japan. The Arboretum and Nursery Office, FFPRI was involved in the maintenance of the *C. japonica* nursery. Ms. Furusawa, FFPRI, performed RNA experiments. Dr. Murai (a retired researcher at FFPRI) conducted artificial crossings, including initiation of inbred lines of *C. japonica*. The sequenced *C. japonica* (‘Kunisaki-3 S3-3’) specimens were housed in the xylarium (TWTw) of FFPRI with voucher (TWTw-29147). We appreciate Miwako Matsumoto, Hisayo Asao, and Dr. Asaka Akita (NIBB) for next generation sequencing.

## Supplementary Materials

**Supplementary Figure 1:**
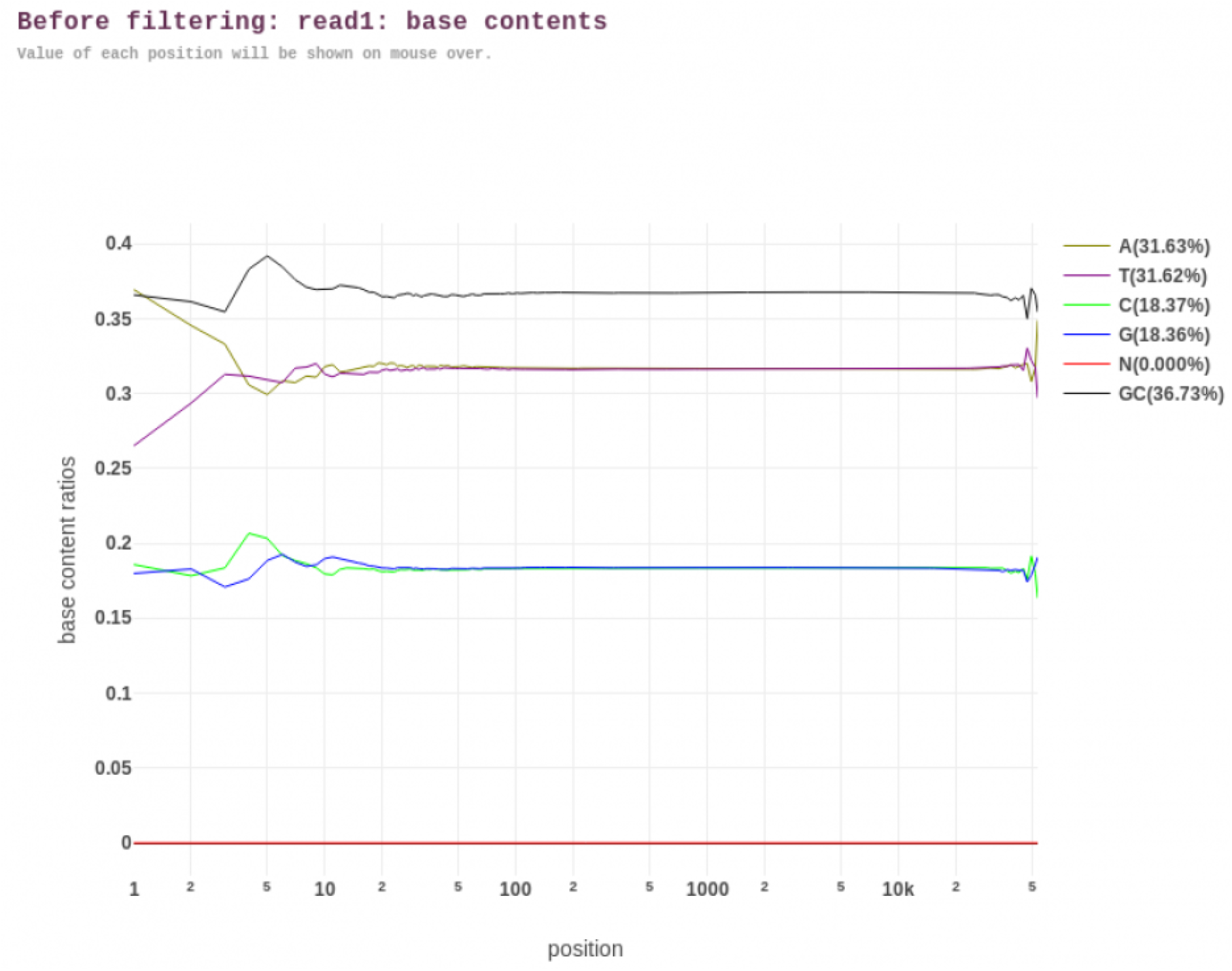
Base Composition Report of the HiFi reads by FastQC (before trimming)

**Supplementary Figure 2:**
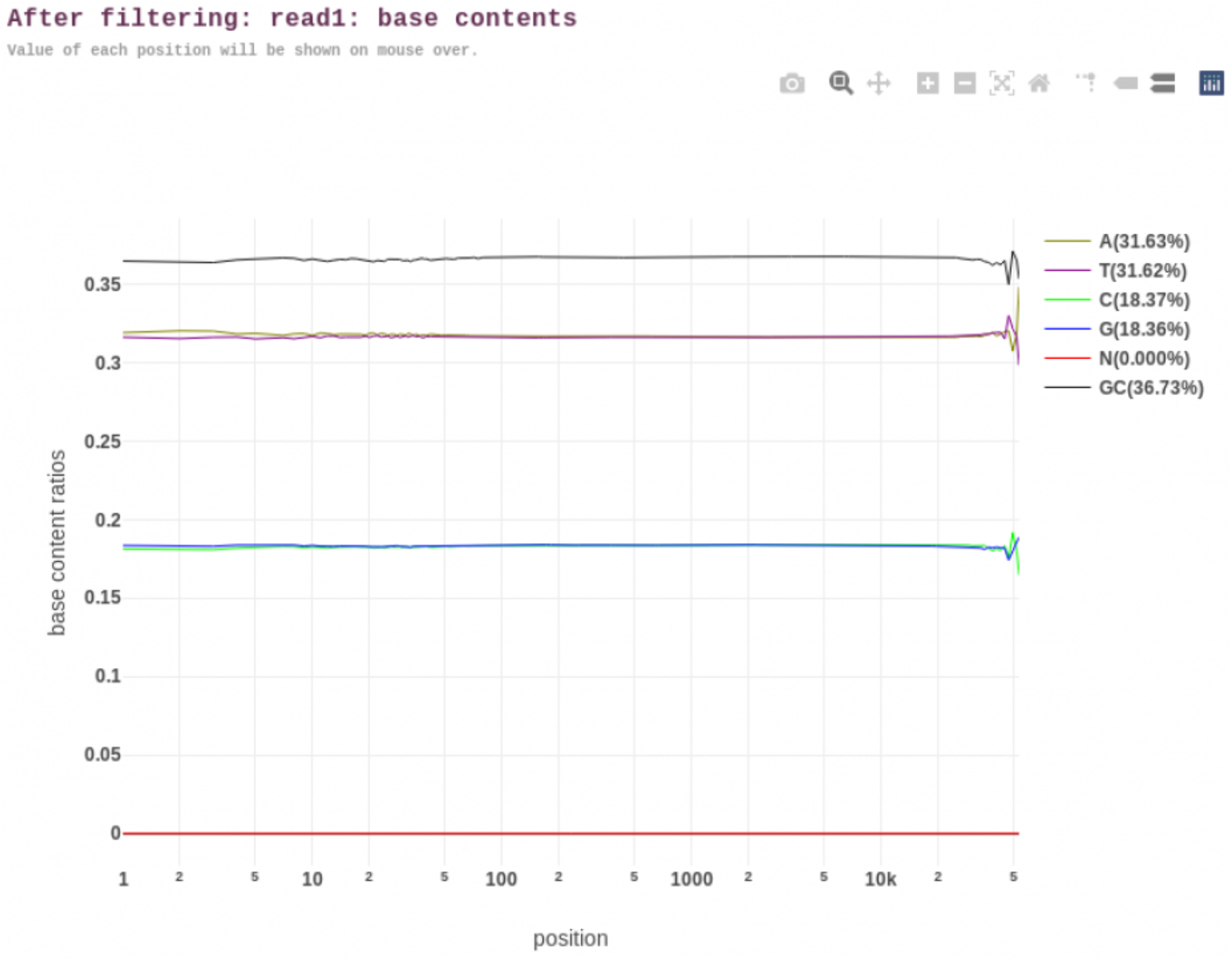
Base Composition Report of the HiFi reads by FastQC (after trimming)

**Supplementary Figure 3:**
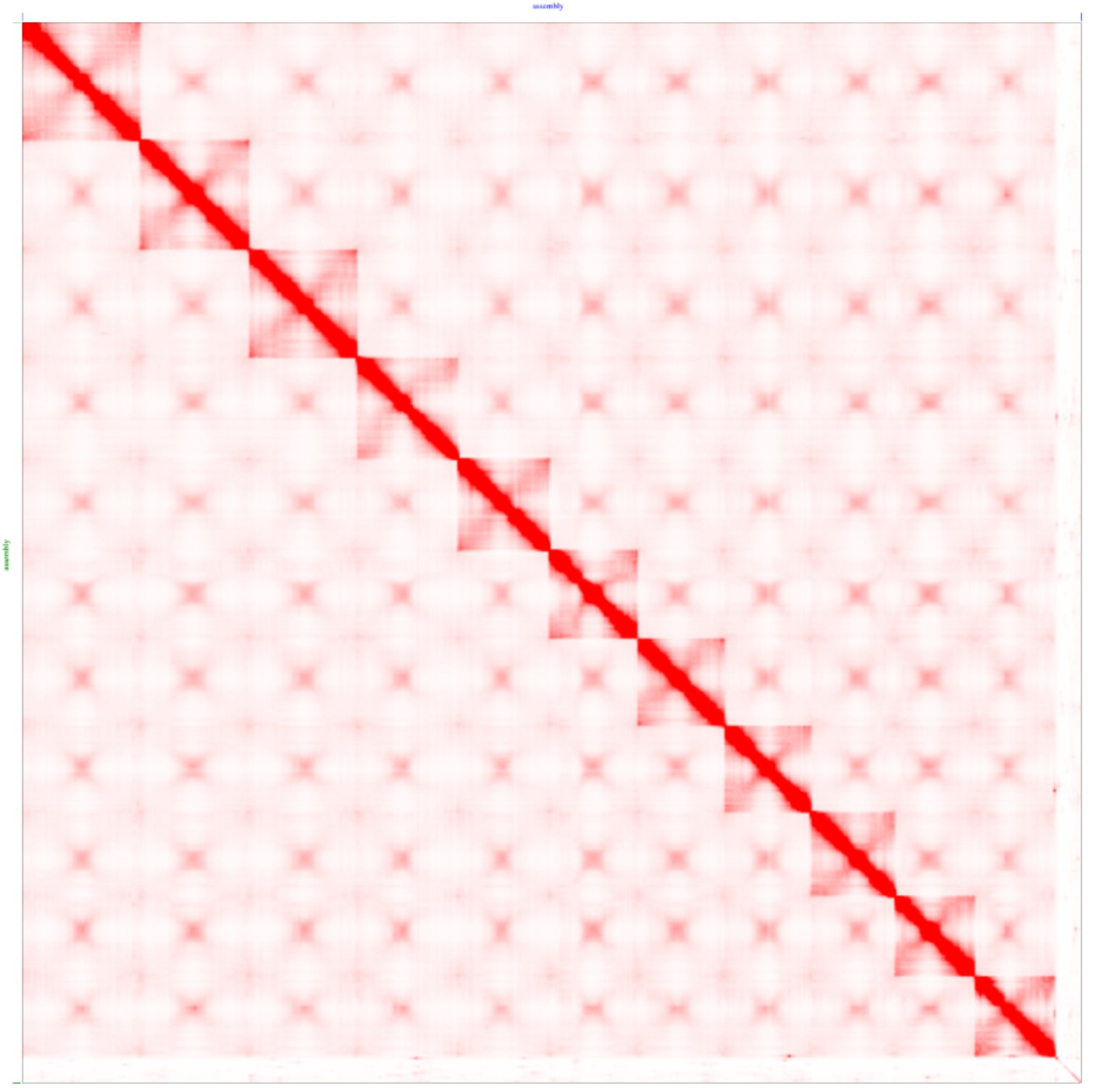
Final Contact Map of the Hi-C analysis (after the manual correction). Eleven chromosomes are clearly seen.

**Supplementary Figure 4:**
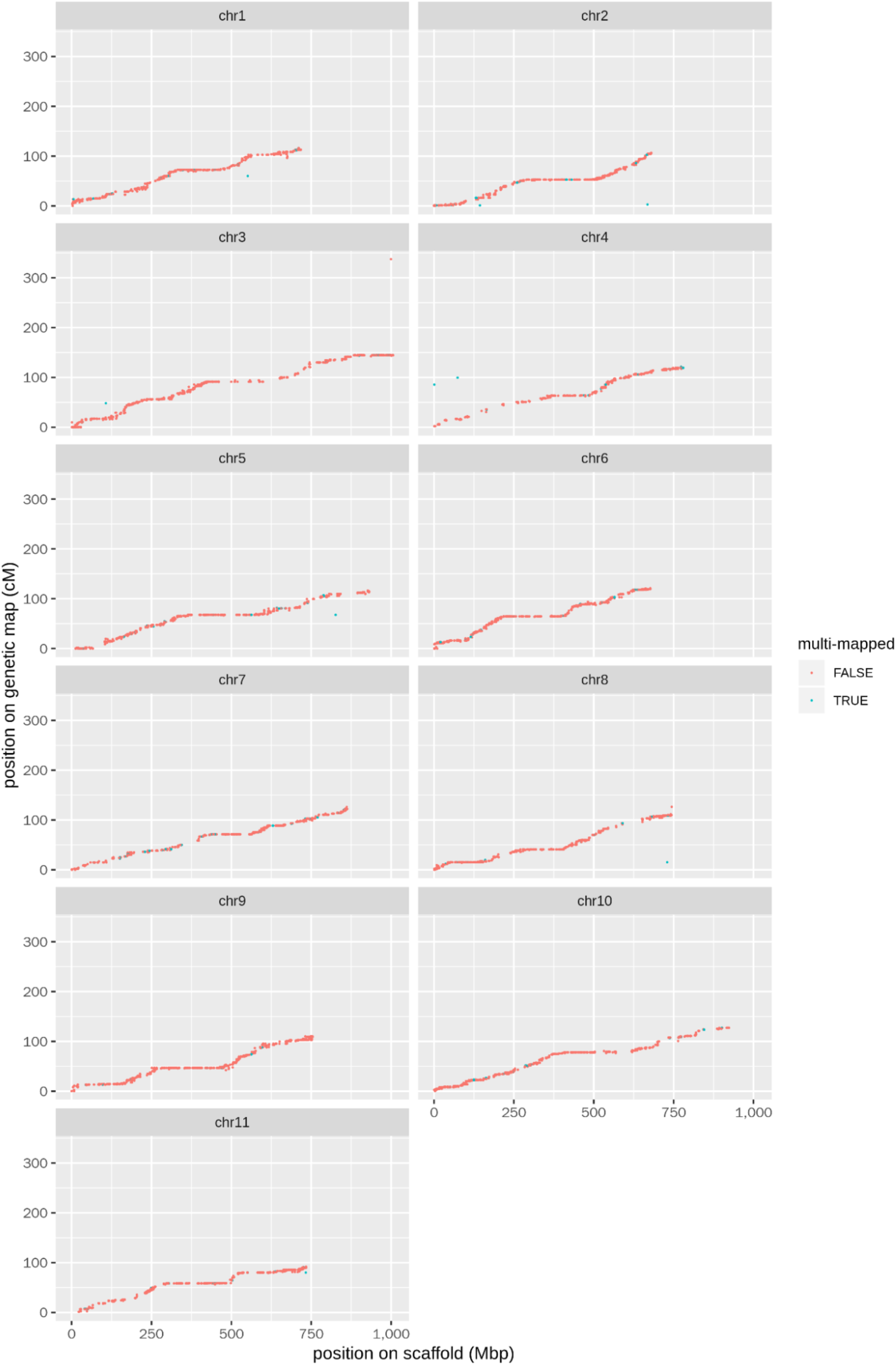
Relationship between the marker positions (cM) and the physical position (Mbp) of the markers. The genetic map and the scaffolds are highly consistent except for the multimapped markers where the sequence aligns to multiple locations on the genome.

